# Memory B cells dominate the early antibody-secreting cell response to SARS-CoV-2 mRNA vaccination in naïve individuals independently of their antibody affinity

**DOI:** 10.1101/2023.12.01.569639

**Authors:** Zhe Li, Anna Obraztsova, Fuwei Shang, Opeyemi Ernest Oludada, Joshua Malapit, Katrin Busch, Monique van Straaten, Erec Stebbins, Rajagopal Murugan, Hedda Wardemann

**Affiliations:** B Cell Immunology, German Cancer Research Center, Heidelberg, Germany; Faculty of Biosciences, University of Heidelberg, Heidelberg, Germany; Cellular Immunology, German Cancer Research Center, Heidelberg, Germany; Faculty of Medicine, University of Heidelberg, Heidelberg, Germany; Structural Biology of Infection and Immunity, German Cancer Research Center, Heidelberg, Germany; Leiden University Center for Infectious Diseases, Leiden University Medical Center, Leiden, The Netherlands

**Keywords:** SARS-CoV-2, mRNA vaccination, affinity maturation, pre-existing memory B cells, somatic hypermutation, germinal center, humoral immunity

## Abstract

Memory B cells (MBCs) formed over the individual’s lifetime constitute nearly half of the adult peripheral blood B cell repertoire in humans. To assess their response to novel antigens, we tracked the origin and followed the differentiation paths of MBCs in the early anti-S response to mRNA vaccination in SARS-CoV-2-naïve individuals on single-cell and monoclonal antibody level. Newly generated and pre-existing MBCs differed in their differentiation paths despite similar levels of SARS-CoV-2 and common corona virus S-reactivity. Pre-existing highly mutated MBCs showed no signs of germinal center re-entry and rapidly developed into mature antibody secreting cells (ASCs). In contrast, newly generated MBCs derived from naïve precursors showed strong signs of antibody affinity maturation before differentiating into ASCs. Thus, although pre-existing human MBCs have an intrinsic propensity to differentiate into ASCs, the quality of the anti-S antibody and MBC response improved through the clonal selection and affinity maturation of naïve precursors.

**Highlights:**

- mRNA vaccination of SARS-CoV-2 naïve individuals recruits naïve and pre-existing MBCs with similar levels of S-reactivity into the response
- S-reactive naïve but not pre-existing MBCs undergo affinity maturation
- S-reactive pre-existing MBCs dominate the early ASC response independent of their antigen affinity
- High-affinity S-reactive MBCs and ASCs develop over time and originate from affinity matured naïve precursors

## Main

A hallmark of the adaptive immune system is the formation of immunological memory. Long-lived humoral immunity is mediated by two types of cells, long-lived antibody-secreting plasma cells that maintain stable antigen-specific serum immunoglobulin (Ig) levels over time and memory B cells (MBCs) that can rapidly differentiate into antibody secreting cells (ASCs) upon re-exposure to the same antigen^1, 2, 3^. Although any recall response will also activate antigen-reactive naïve B cells that encounter the antigen for the first time, their affinity is usually lower than that of MBCs that have undergone extensive affinity maturation in germinal center (GC) reactions during the primary response^1^. Consequently, pre-existing antigen-specific MBCs dominate humoral recall responses initiated by the same pathogen. The fast production of MBC-derived high-affinity serum antibodies mediates immediate and effective protection from infections with pathogens that show little antigenic variation. However, antigenic variation among pathogen populations frequently allows pathogens to evade MBC-derived humoral immunity. In re-infections with variant influenza strains, high-affinity MBCs suppress *de novo* responses from naïve precursors, an effect described as original antigenic sin or more adequately as antigenic, immunogenic or immune imprinting^4, 5^.

In humans, class-switched and non-class-switched MBCs constitute nearly half of the circulating B cell pool^6^. All subsets express mutated Ig genes with clear signs of antigen-mediated selection suggesting that the cells show high affinity for foreign antigens that the host encountered earlier in life. Because MBCs develop throughout ongoing immune responses, their antigen-receptor repertoire shows a broad range of affinities likely enabling flexibility in recall responses to antigenically distant stimuli^7^. Indeed, MBCs with reactivity to seasonal human coronaviruses (HCoVs) participate in infection or vaccination-induced anti-SARS-CoV-2 responses if their antigen receptors show strong cross-reactivity with the viral spike (S) proteins, predominantly the conserved S2 subunit^8, 9^. Direct lymph node sampling showed that cross-reactive MBCs can participate in GC reactions along with naïve B cells^10, 11^. However, to what extent pre-existing MBCs participate in the anti-SARS-CoV-2 response compared to naïve B cells and affinity mature over time in individuals with no prior exposure to the virus is not well understood.

To address this question, we studied the evolution of the anti-S protein response in SARS-CoV-2 naïve individuals over two vaccinations with mRNA. Despite similar levels of S-reactivity at the onset of the response compared to naïve B cells, pre-existing S-reactive MBCs differentiated into ASCs. In contrast, newly developing MBCs derived from activated S-reactive naïve B cells showed evidence of efficient affinity maturation in GC reactions upon class-switching to IgG. The data demonstrate the intrinsic propensity of MBCs to drive immediate humoral immune responses independently of their antigen-binding strength through differentiation into ASCs. Although rare pre-existing MBCs may participate in GC reactions, the quality of the anti-S IgG memory antibody response to mRNA vaccination is based on the antigen-mediated activation and affinity maturation of naïve B cells likely during GC reactions.

## Results

### The early B cell response to mRNA vaccination in SARS-CoV-2 naïve individuals

To follow the development of the anti-S B cell response upon mRNA vaccination, we collected plasma and peripheral blood mononuclear cells (PBMCs) from four healthy SARS-CoV-2-naïve volunteers (V1-4) directly before (pre) and one, two, and three weeks (1w, 2w, 3w) after the first (I) and second (II) vaccination with Comirnaty® (Fig. 1a). All donors responded with strong humoral IgG but not IgM or IgA responses against the viral S protein, especially after the boost (Extended Data Fig. 1a-c). The lack of IgA titers and delayed IgG response compared to an individual with prior coronavirus disease (V5) confirmed that these donors were SARS-CoV-2 naïve. Indicative of the ongoing immune response, flow cytometric analyses showed an increase in S+ naïve B cells (CD19+CD27-S+) and MBCs (CD19+CD27+S+) after the prime and boost vaccinations and of ASCs (CD19+CD27+CD38+) predominantly after the prime (Fig. 1b,c).

**Fig. 1:**
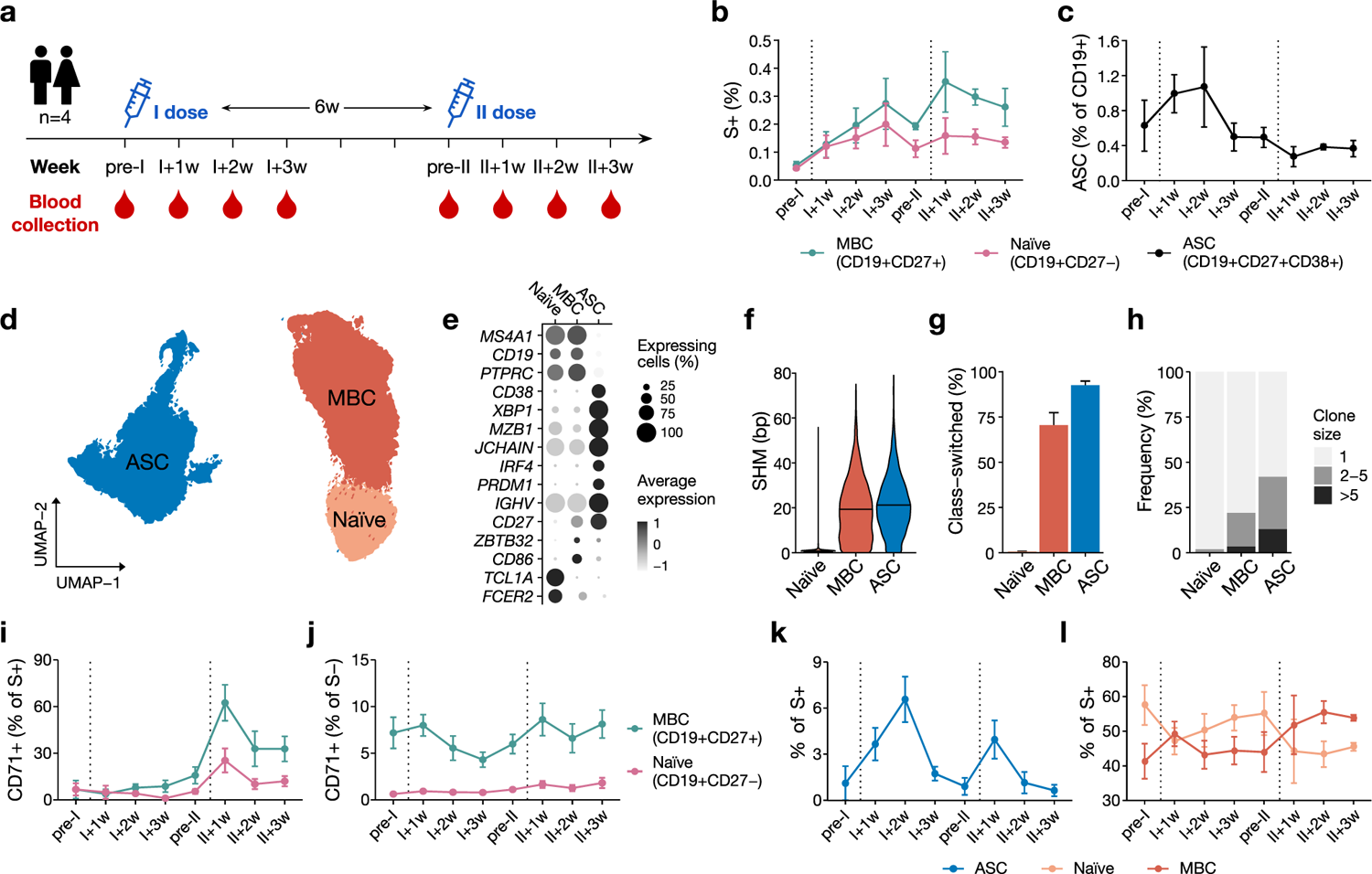
Longitudinal characterization of early B cell response against SARS-CoV-2 mRNA vaccination. **a**, Study design. I – first dose; II – second dose; pre-I and pre-II – the day before the first and the second vaccination; 1w, 2w, 3w – one, two and three weeks after vaccination. **b**, Frequency of S+ B cells in memory (CD19+CD27+) and naïve (CD19+CD27-) population over time. **c**, Frequency of ASCs over time. **d**, UMAP projection of peripheral blood B cells transcriptional profiles colored by population defined by transcriptome analysis. ASC – antibody-secreting cell; MBC – memory B cell. **e**, Expression profile of selected marker genes. **f,** *IGHV*+*IGK/LV* somatic hypermutation (SHM) counts across the cell populations. **g**, Class-switched antibody frequency. **h**, Clone size distribution. **i-j**, Frequency of CD71+ cells in S+ (**i**) and S-(**j**) naïve (CD19+CD27-) and memory B cells (CD19+CD27+). **k-l**, Frequency of antibody secreting (**k**) and B cell (**l**) populations in S+ cells over time. Dots show average frequency across the volunteers and error bars show SEM. Dotted vertical lines indicate prime and boost.

To characterize the response more deeply, we sorted these populations using barcoded S baits and performed transcriptome and Ig gene analyses (Fig. 1b-h). We included the activation marker CD71 and also isolated S-CD71+ naïve and MBCs to capture recently activated cells whose bait-reactivity was below the flow cytometric detection threshold^12^ (Extended Data Fig. 1d). The transcriptome and Ig gene analysis confirmed the cellular identity of the sorted populations (Fig. 1e-h). ASCs were identified by high expression of *CD38, IRF4* and *IGHV* and highly mutated and class-switched Ig genes and showed clear signs of clonal expansion (Fig. 1e-h). MBCs were overall more similar to naïve B cells in their expression profile but expressed *CD27*, the activation marker *CD86*, and showed signs of clonal expansion (Fig. 1e,h). As expected, their Ig genes were highly mutated and predominantly class-switched (Fig. 1f,g). The frequency of S+ CD71+ naïve and MBCs increased with time, especially after the boost compared to little change in the S-populations (Fig. 1i,j). Independently of their S-reactivity, CD71+ cells were more frequent among MBCs than naïve B cells, indicative of a higher baseline activation status (Fig. 1j). The transcriptome analysis confirmed the higher expression of *TRFC* encoding for CD71 in MBCs compared to naïve B cells, although only the S+ *TRFC*+ subset reflected the vaccination induced changes detected by flow cytometry (Extended Data Fig. 1e). Because S-CD71+ naïve and MBCs showed no evidence of active participation in the vaccine response and might therefore reflect bystander activation, we focused all further analyses on S+ cells.

S+ ASCs were overall rare, presumably due to their low surface BCR expression. Nevertheless, their frequency increased after each vaccination (Fig. 1k). Before and after the first dose, naïve B cells constituted the majority of S+ cells, whereas MBCs dominated the response after the boost, likely reflecting the ongoing differentiation of antigen-reactive naïve precursors into MBCs (Fig. 1l).

Collectively, the data show that mRNA vaccination recruited not only S-reactive naïve B cells into the response but also highly mutated pre-existing S-reactive MBCs and induced the development of S+ and S-ASCs along with strong humoral anti-S IgG responses after the boost.

### Low-but not high-SHM IgG MBCs show signs of active GC participation

To dissect the contribution of newly generated and pre-existing MBCs to the anti-S response in more detail, we tracked the mutation load of S+ IgG MBCs. At I+1w, the vast majority (95%) of S+ IgG MBCs were highly mutated comparable to the pre-vaccination levels (Fig. 2a). Only few cells were unmutated or lowly mutated at this early time point, but their numbers increased by II+1w suggesting that newly recruited naïve B cells had class-switched to IgG and entered the MBC pool (Fig. 2b). To assess the contribution of these two S+ IgG MBC subsets to the response over time, we separated S+ IgG MBCs with no or low-SHM counts from those with high-SHM counts (referred to as low-SHM MBCs and high-SHM MBCs, respectively) based on the near bimodal distribution of their SHM load (Fig. 2a and Extended Data Fig. 2a). Low-SHM IgG MBCs accumulated SHM with time, likely reflecting their export from ongoing GC reactions (Fig. 2c) and suggesting that naïve B cells differentiate into MBCs and class-switched to IgG before hypermutating their Ig genes. In contrast, lowly mutated IgM MBCs were detected with similar frequency at all time points and showed no change in SHM load over time indicating that these cells did not undergo Ig gene diversification in response to the vaccination (Extended Data Fig. 2b-d).

**Fig. 2:**
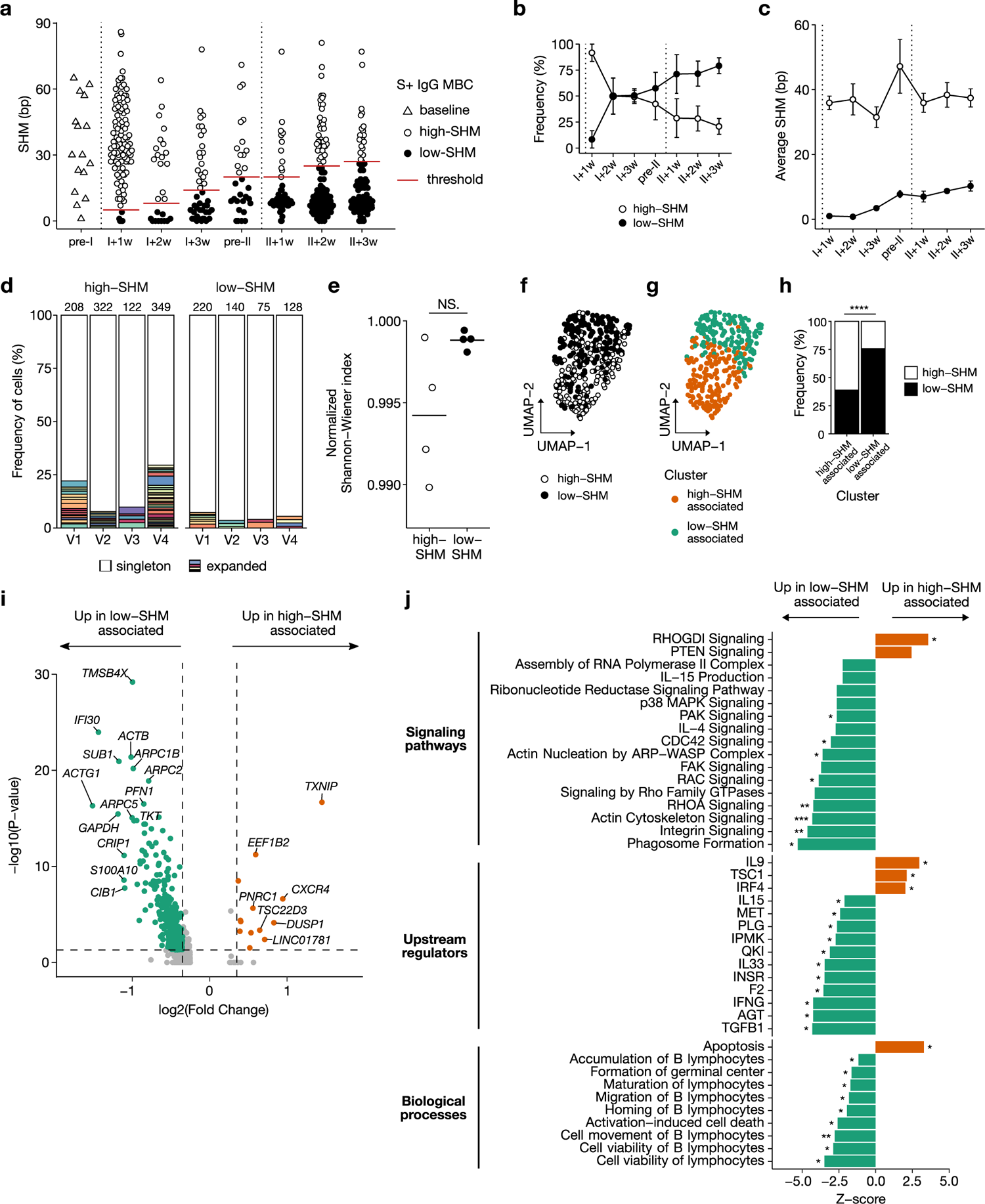
Low-SHM but not high-SHM S+ IgG MBCs display signs of ongoing GC reaction. **a,** *IGHV + IGK/LV* SHM counts in S+ IgG MBCs. Dotted vertical lines indicate prime and boost. **b,** Frequency of low- and high-SHM MBCs over time. **c,** Average SHM count over time. **d,** Clonal expansion of low- and high-SHM MBCs. **e**, Clonal diversity. Each dot represents one donor. **f-g,** UMAP projection of high- and low-SHM S+ IgG MBCs colored by SHM level (**f**) or cluster defined by unsupervised transcriptome clustering (**g**). **h**, Frequency of high- and low-SHM cells in the transcriptome clusters. **I,** Genes differentially expressed between low- and high-SHM associated clusters. (**j**) Signalling pathways and transcriptional programs enriched in genes differentially expressed between low- and high-SHM associated clusters. NS. - not significant, *P < 0.05, **P < 0.01, ***P < 0.001, ****P < 0.0001, Pearson’s Chi-squared test with Yates’ continuity correction (**h**), Fisher’s exact test (**j**). Error bars display SEM across donors.

The average SHM count in high-SHM IgG MBCs remained stable even within large persistent clones (Fig. 2c and Extended Data Fig. 2e) suggesting that these cells originated from different precursors than the low-SHM IgG subpopulation. Indeed, we observed little clonal overlap (n=7) between expanded B cell clones in low-SHM (n=36) and high-SHM (n=108) MBCs, and we identified higher numbers of expanded B cell clones with longer tree branches reflecting a longer Ig gene diversification history and lower clonal diversity in high-SHM compared to low-SHM MBCs (Fig. 2d,e and Extended Data Fig. 2f,g). The two populations also differed in their transcription profiles (Fig. 2f-h). Upregulated genes and signaling pathways linked to low-SHM MBCs were associated with cytoskeleton remodeling (*ACTB, ACTG1, PFN1, ARPC2* and *ARPC1B)*, cell motility, adhesion and proliferation (RhoA, Cdc42, Rac, Pak, Fak, Actin, Integrin signaling) as well as T cell help, B cell activation, GC formation and viability (IL-4, p38 MAPK, IL-15 and IFNγ signalling) suggesting that these cells are recent GC emigrants^13, 14^ (Fig. 2i,j). In contrast, upregulated genes associated with high-SHM MBCs were mostly linked to plasma cell differentiation and apoptosis (*TXNIP*, IRF4 and IL-9 signalling)^15, 16^.

We conclude that S+ low-SHM IgG MBCs develop from naïve precursors that undergo Ig gene diversification and accumulate in the S+ IgG MBC pool over time. These cells are not related to pre-existing S+ IgG MBCs with high-SHM counts, which likely developed prior to the vaccination, lack signs of ongoing affinity maturation, and express genes associated with plasma cell differentiation.

### Naïve B cell-derived but not pre-existing MBCs undergo anti-S affinity maturation

Next, we determined whether high- and low-SHM IgG MBCs differed in their S reactivity. Low-SHM IgG MBCs showed on average higher S and RBD reactivity and stronger signs of BCR activation than high-SHM MBCs based on barcode read counts and signaling pathway gene expression scores, respectively (Fig. 3a,b). Both populations also differed in their paired V gene usage (Fig. 3c). Low-SHM MBCs showed a strong enrichment of *IGHV, IGKV,* and *IGLV* genes and Ig gene features associated with S reactivity (Fig. 3d,e) including antibodies with long HCDR3^17, 18^ and *IGHV3-53* or *IGHV3-66* segments paired with *IGHJ6* that frequently encode high-affinity anti-RBD antibodies^19^ (Fig. 3f,g).

**Fig. 3:**
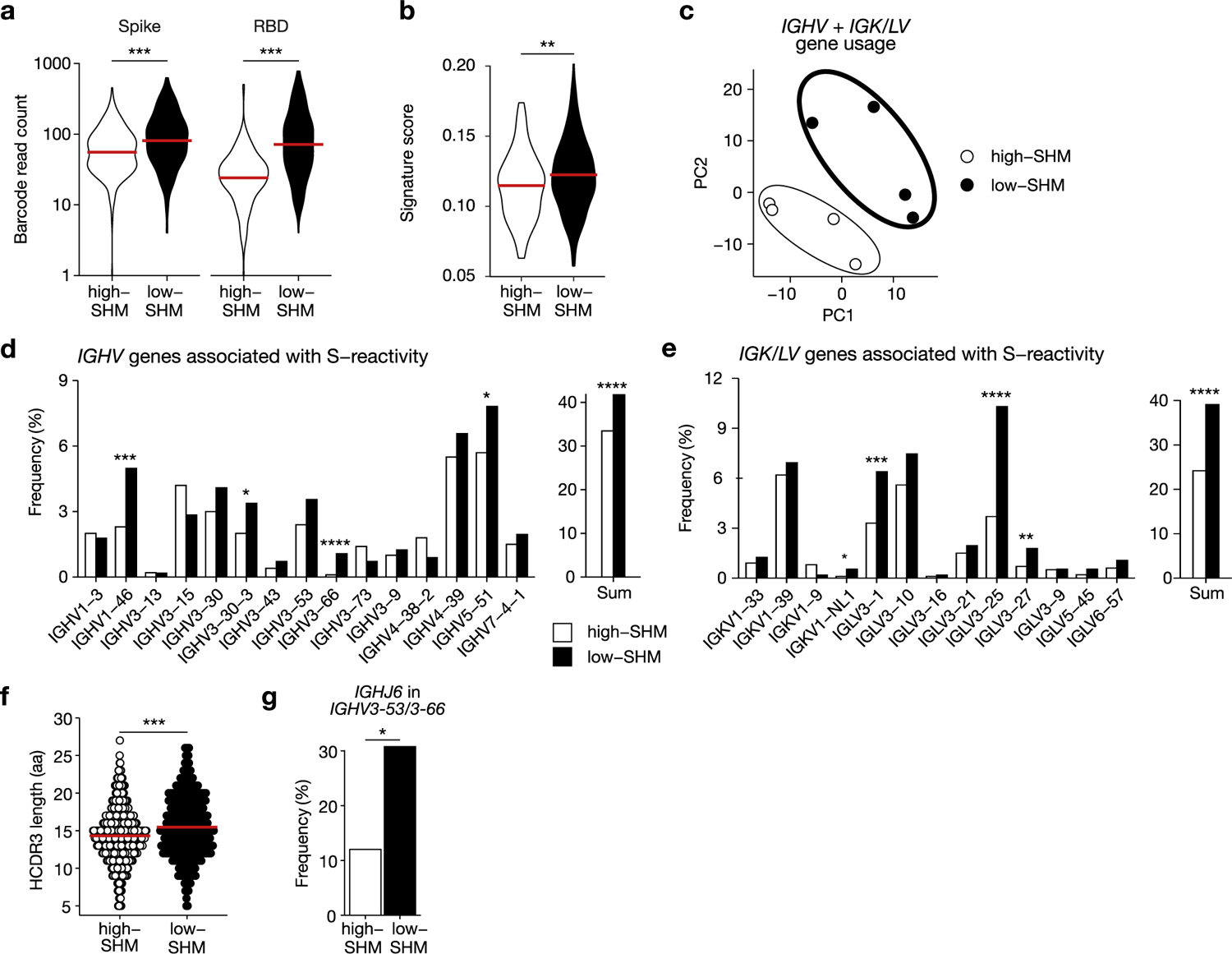
Features associated with S-binding are enriched in antibodies of low-SHM S+ MBCs. **a**, Read counts of Spike and RBD tetramer barcodes in S+ IgG MBCs. **b**, BCR signaling pathway gene expression in S+ IgG MBCs. **c-f**, Ig variable segment gene usage of S+ MBCs. **c**, Principal component analysis (PCA) on paired chain V gene usage (dots represent MBC populations from individual donors). **d**, Frequency of individual *IGHV* genes associated with S-binding (left) or their pooled frequency (right). **e**, Frequency of individual *IGK/LV* genes associated with S-binding (left) or their pooled frequency (right). **f**, HCDR3 length distribution in S+ MBCs. **g**, Frequency of *IGHJ6* usage in cells using *IGHV3-53* or *IGHV3-66* in S+ MBCs. *P < 0.05, **P < 0.01, ***P < 0.001, ****P < 0.0001, two-tailed Mann-Whitney test (**a**-**b** and **f**), exact binomial test with Benjamini-Hochberg correction for multiple testing (**d**-**e** and **g**).

To evaluate the binding strength of low- and high-SHM MBCs, we cloned and expressed 141 monoclonal antibodies (mAbs) from S+ IgG MBCs and 25 mAbs from S+ naïve B cells isolated at different time points after the vaccination (Supplementary Table 1 and Extended Data Fig. 3a,b). Already after the prime, mAbs from low-SHM IgG MBCs showed on average higher antigen binding levels compared to mAbs from high-SHM IgG MBCs (Fig. 4a). Their binding strength increased after the boost, whereas that of high-SHM IgG MBCs remained similar to naïve B cells (Fig. 4a,b). Affinity measurements of S-reactive mAbs with RBD specificity confirmed that low-SHM IgG MBC mAbs showed overall better antigen-reactivity than mAbs from high-SHM MBCs and that their binding strength and SHM count correlated positively, likely as a result of active affinity maturation in GC reactions (Fig. 4c and Extended Data Fig. 3c,d). ELISAs with S protein from four of the most prevalent HCoVs (HKU1, OC43, NL63, 229E) showed no difference in reactivity between mAbs from low- and high-SHM IgG MBCs after the prime or boost and only a few mAbs showed high HCoVs cross-reactivity suggesting that HCoVs cross-reactivity played no major role in the recruitment of low- or high-SHM IgG MBCs into the response and that highly cross-reactive MBCs were overall rare (Fig. 4d). The finding is in agreement with the lack of measurable anti-HCoV serum IgG antibody levels after both vaccinations (Extended Data Fig. 3e).

**Fig. 4:**
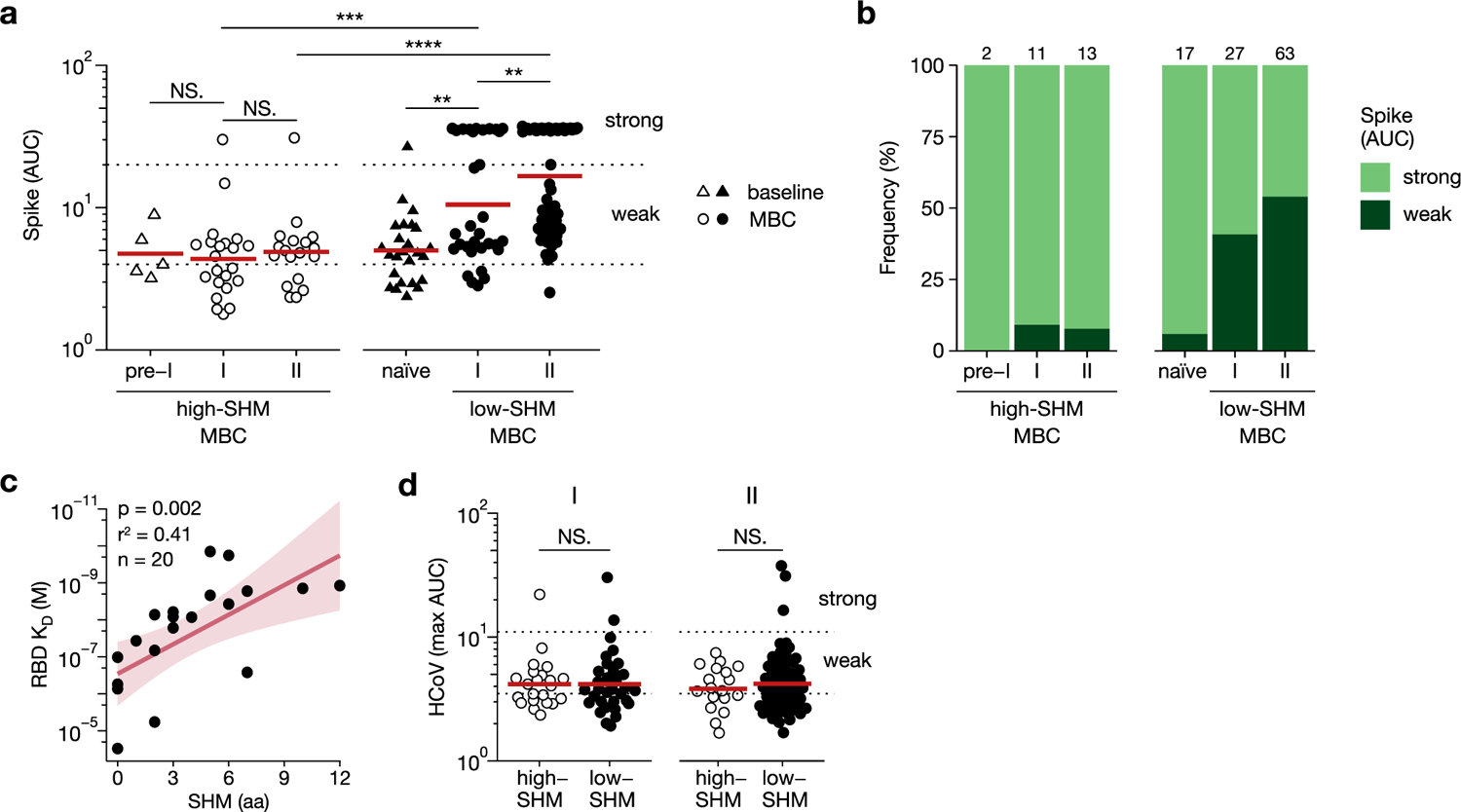
Antibodies of low-SHM IgG MBCs are better S binders that are improving over time. **a-b**, ELISA AUC values (**a**) and frequencies of strong and weak S binders (**b**) in low- and high-SHM S+ IgG MBCs separated by isolation time point (I – after first vaccination, II – after second vaccination) and in naïve B cells. Red lines show average, dotted horizontal lines show thresholds for weak- and strong-binding mAbs. Numbers on top of the bars show the numbers of antibodies expressed and tested. **c**, RBD binding affinity of strongly RBD binding low-SHM IgG mAbs versus total *IGHV* + *IGKV/IGLV* SHM counts. Data are representative of at least two independent experiments. **d**, HCoV Spike ELISA AUC values of mAbs from high- and low-SHM MBCs. HCoV AUC indicates the highest value obtained for each mAb among four tested antigens (HKU1, OC43, NL63, 229E). *P < 0.05, **P < 0.01, ***P < 0.001, ****P < 0.0001, NS, not significant; two tailed Mann-Whitney test.

Thus, high binding anti-SARS-CoV-2 S mAbs developed predominantly from low-SHM IgG MBCs by affinity maturation, whereas high-SHM IgG MBCs mAbs showed overall low antigen reactivity, which did not improve over time or after the booster vaccination.

### Predominant differentiation of high-SHM IgG MBCs into ASCs despite weak anti-S antibody reactivity

To determine to what degree low- and high-SHM MBCs differentiated into ASCs over the course of the response, we examined the Ig gene repertoire of low- and high-SHM ASCs (Extended Data Fig. 4a). Paired Ig heavy and light chain V gene usage and clonal overlap analyses provided evidence for the common origin of ASCs and MBCs with low-SHM count and of ASCs and MBCs with high-SHM load, respectively (Fig. 5a,b). High-SHM IgG ASCs dominated the response at all time points, although the frequency of low-SHM ASCs increased after each vaccination (Fig. 5c). mAbs cloned from low-SHM ASCs showed stronger S-reactivity and the frequency of antigen-binders was overall higher in low-compared to high-SHM ASCs, similar to the differences we observed between low- and high-SHM IgG MBCs (Fig. 5d). In line with these findings, low-SHM ASCs showed higher S-barcode read counts and S+ cell frequency compared to their high-SHM counterparts (Extended Data Fig. 4b,c).

**Fig. 5:**
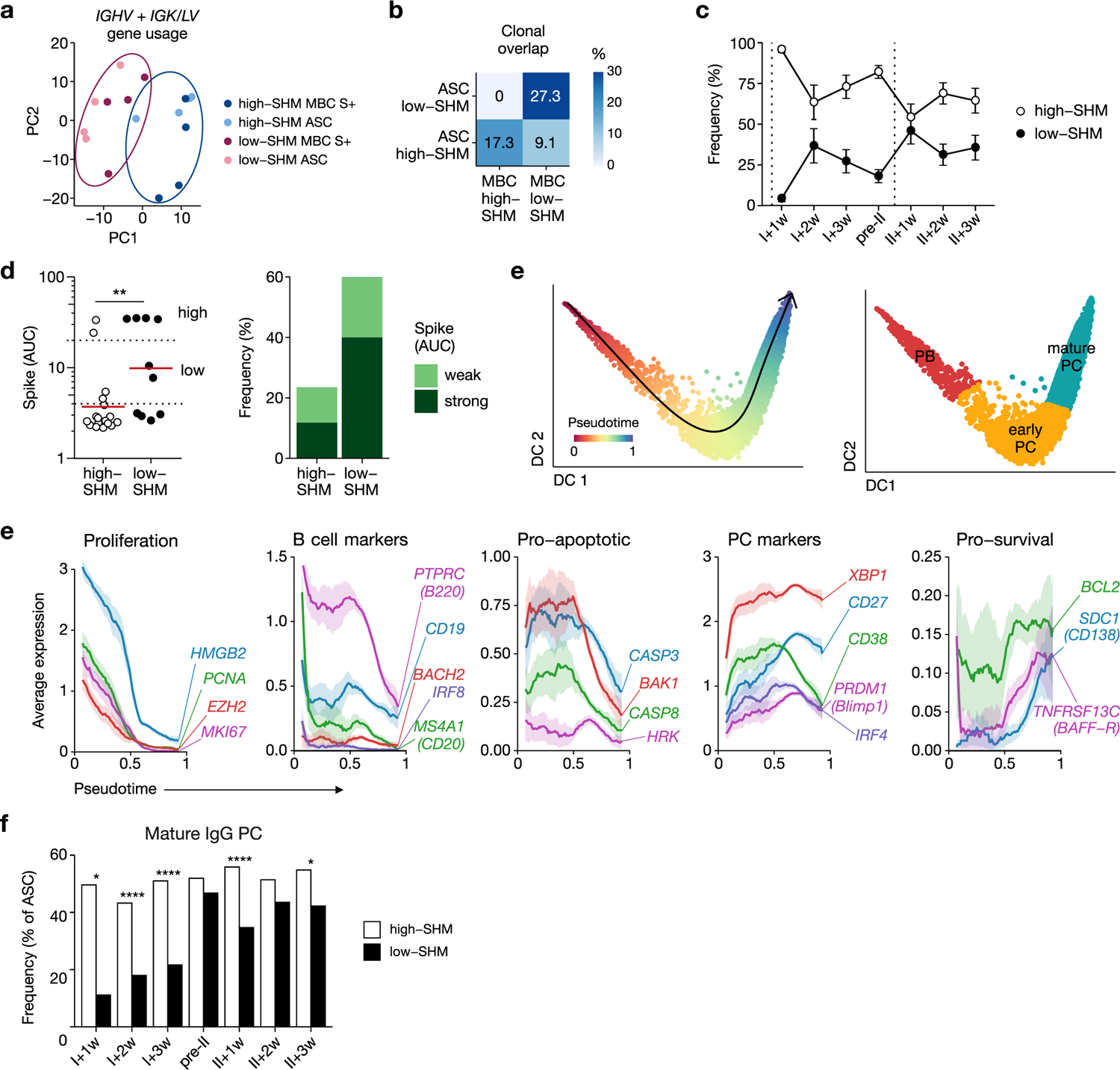
Naïve B cells generate less mature but more potent IgG expressing ASCs compared to pre-existing MBCs. **a**, Principal component analysis (PCA) on paired chain V gene usage of IgM and IgG cells. Dots represent MBCs or ASCs populations from individual donors. **b**, Clonal overlap between IgM or IgG high- and low-SHM S+ MBCs and ASCs. **c**, Frequency of high- and low-SHM IgG ASCs over time. Dotted vertical lines indicate prime and boost. **d**, ELISA AUC values (left) and frequencies of S-binding mAbs (right) in ASCs clonally related to S+ MBCs. Dotted lines show thresholds for weak- and strong-binding mAbs. Numbers on top of the bars show the numbers of antibodies expressed and tested. Red lines show average. **e**, Diffusion map of ASCs colored by pseudotime (left) or maturation stage (right). PB, plasmablasts (pseudotime ⩽0.33); early PC, early plasma cell (0.33 < pseudotime < 0.66); mature PC, mature plasma cell (pseudotime ⩾0.66). **f**, Expression of genes contributing in mature plasma cell phenotype along the pseudotime trajectory. **g**, Distribution of ASC maturation stages between low- and high-SHM IgG ASCs across time points. *P < 0.05, **P < 0.01, ***P < 0.001, ****P < 0.0001, binomial test (**g**), two tailed Mann-Whitney test (**d**).

To be able to link antigen-reactivity to ASC differentiation kinetics, we applied pseudotime analysis based on diffusion distance (Fig. 5e). The expression of proliferation markers, B cell markers, and pro-apoptotic genes decreased gradually along the pseudotime trajectory, while plasma cell and survival markers, including CD138, increased concomitantly (Fig. 5f). Thus, the pseudotime values reflected the ASC maturation from proliferating plasmablasts (PB) to mature plasma cells (PC) over time and was used to define the maturation stages of ASCs with high- or low-SHM counts (PB, early PC and mature PC, Fig. 5e). Already at I+1w, high-SHM IgG ASCs had higher pseudotime values compared to low-SHM IgG ASCs with high and stable frequency of mature PCs over time (Fig. 5g and Extended Data Fig. 4d). In contrast, mature PCs were rare among low-SHM IgG ASCs after the first vaccination and increased only slowly mainly after the boost concomitant with increase in anti-S serum IgG titers (Fig. 5g and Extended Data Fig. 1a).

Thus, in response to the vaccination, S+ high-SHM IgG MBCs expressing weakly binding anti-bodies developed quickly into non-proliferating mature ASCs. In contrast, low-SHM MBCs with similar antigen reactivity improved their antibody affinity by SHM in GC reactions before differentiating into proliferating ASCs (Extended Data Fig. 4g).

## Discussion

Our data demonstrate that the initial anti-S ASC response to mRNA vaccination in SARS-CoV-2 naïve individuals is driven by pre-existing MBCs although the average S-binding strength of their antibodies was not higher than that of newly developing MBCs derived from naïve precursors. We have made similar observations in malaria-naïve individuals after vaccination with live parasites suggesting that human MBCs have a high propensity to differentiate into ASCs upon BCR-mediated activation even when they have not previously encountered the stimulating antigen^20^. The high numbers of SHM in the S+ IgG MBCs that responded to SARS-CoV-2 vaccination in our study were comparable to pre-vaccination levels indicating that these cells must have developed prior to the vaccination. Their relatively low SARS-CoV-2 S-protein reactivity likely reflects weak antigen cross-reactivity, a common feature of Ig molecules including antibodies that have undergone affinity maturation^21^. Given the size of the human MBC pool, cross-reactivity in this compartment likely plays an important role in protection from diverse antigenic threats by increasing the number of cells that can be recruited into the response and contribute to the rapid production of serum antibodies by differentiation into ASCs. The low anti-S antibody titers after the prime likely reflect the weak binding properties of these cells compared to the strong binders that develop after the boost from affinity-matured naïve precursors.

Numerous studies illustrated that human MBCs readily respond to antigen-independent polyclonal stimuli and bystander T cell help by proliferation and differentiation into ASCs *in vitro* ^22, 23^. Especially IgG class-switched MBCs have been shown to have a lower activation threshold and higher sensitivity to bystander T cell help than IgM MBCs^24, 25, 26^. The high-SHM MBCs studied here displayed similar levels of S-reactivity compared to the low-SHM MBCs, nevertheless the two subsets differed in their differentiation paths. The strong T cell responses induced by mRNA vaccination might have promoted the fast differentiation of S-reactive IgG MBCs into ASCs compared to naïve B cells, which showed clear signs of affinity maturation in GC reaction^1, 27, 28^.

Likely due to the highly polyclonal nature of the anti-S response, we did not observe strong signs of clonal expansion among pre-existing high-SHM MBCs over the vaccination course. Instead, the cells upregulated TXNIP, IL-9 and IRF4 expression (markers associated with PC differentiation, at least in mice)^15, 16^ and rapidly differentiated into more terminally differentiated ASCs. The fast development of pre-existing MBCs into ASCs suggests that these cells are the main contributors to the first wave of serum IgG. The fast secretion of low-binding anti-S IgG antibodies by these cells might promote the activation and selection of S-reactive B cells through the formation of immune complexes and subsequent deposition of antigen on follicular dendritic cells GC^29^. IgA MBCs may play a similar role since highly mutated IgA-secreting cells constituted a significant fraction of ASCs, but serum anti-S IgA was low (Extended Data Fig. 4e,f).

The second vaccination induced a strong increase in titers and likely antibody quality, since we did not detect a proportionally strong increase in the frequency of ASCs after the boost. Our data suggest that high-affinity serum IgG antibodies originated predominantly from de novo responses of naïve B cell precursors, which class-switched and showed clear signs of affinity maturation prior to differentiation into proliferating ASCs. Although it is possible that some pre-existing MBCs in our study participated in GC reactions, these were likely rare events relative to the GC participation of naïve B cell clones including cells expressing Ig genes associated with S-reactivity^30^. Direct sampling of lymph node GC cells in SARS-CoV-2 vaccinees identified highly mutated clones with strong cross-reactivity to S-protein from HCoVs suggesting that these cells represented pre-existing MBCs that had developed in response to prior infections with common cold viruses^11^. Nevertheless, the majority of GC B cells in this study seemed to be SARS-CoV-2 S-specific and likely developed from naïve precursors since their mutation load was significantly lower than that of clones with HCoVs cross-reactivity.

In summary, our study demonstrates that human IgG MBCs have an intrinsic propensity to differentiate into ASCs and the quality of the response to mRNA vaccination in SARS-CoV-2 naïve individuals improves through the clonal selection and affinity maturation of potent S-reactive naïve B cell precursors.

## Methods

### Study design, sample collection and storage

This study was approved by the ethics committee of the medical faculty of the University of Heidelberg (S-0001-2022). Written consent was obtained from all participants. The study was conducted according to the principles of the Declaration of Helsinki and also conformed to the principles set out in the Department of Health and Human Services Belmont Report. Five healthy donors were enrolled and received two doses of BNT162b2 mRNA SARS-CoV-2 vaccine (Comirnaty®) that encodes a prefusion stabilized, membrane-anchored SARS-CoV-2 full-length Spike protein. Peripheral blood samples were collected one day before, one, two and three weeks after each of the two vaccinations. Peripheral blood mononuclear cells (PBMCs) were isolated using Ficoll gradient density centrifugation, frozen in 10% dimethyl sulfoxide in fetal calf serum and stored in liquid nitrogen until further use.

### Antigens

The mammalian expression vector encoding SARS-CoV-2 Spike protein or RBD derived from the first virus isolate, Wuhan-Hu-1 was previously described^31^. Two modifications were introduced to Spike protein sequences to stabilize the trimer in the pre-fusion conformation^31^. Spike and RBD were expressed in HEK293F cells and purified by NI-IMAC resin. For constructing baits for flow cytometry, purified SARS-CoV-2 Spike and RBD were biotinylated using the EZ-Link NHS-Biotin (Thermo Fisher Scientific) following the manufacturer’s instructions. The mole-to-mole ratio of biotin to protein was quantified using the Pierce Biotin Quantitation Kit (Thermo Fisher Scientific). Biotinylated Spike was conjugated to streptavidin-FITC (BioLegend) and streptavidin-PE (BioLegend, TotalSeq™-C0951) and RBD to streptavidin-APC (BioLegend, TotalSeq™-C0956) overnight at 4°C.

### Flow cytometry, cell sorting and 10X sample preparation

PBMCs were thawed at 37°C and washed twice with RPMI (Gibco). Cells were incubated for 10 min at 4 °C with Fc receptor block (TruStain FcX, BioLegend, 1:10). Cells were washed and incubated in FACS buffer (1xPBS, 2%FCS) with the following anti-human antibodies: CD19-BV785 (BioLegend, 1:10), CD3-APC-eFluor 780 (Invitrogen, 1:200), CD8-APC-eFluor 780 (Invitrogen, 1:200), CD14-APC-eFluor 780 (Invitrogen, 1:200), CD16-APC-eFluor 780 (Invitrogen, 1:200), CD27-PE-Cy7 (BioLegend, 1:20), CD38-BV605 (BD Biosciences, 1:20), CD71-BV510 (BD Biosciences, 1:10), TotalSeq-C hashtag antibodies 1–12 (BioLegend, 1:100) and Spike and RBD antigen tetramers. Afterwards, cells were washed and stained with 7-Aminoactinomycin D (7AAD) (Invitrogen, 1:400) that was used as dead cell marker. Single cells were sorted with FACSAria III (BD Biosciences) into cooled 1.5-ml tubes. FACS data were collected with the BD FACSDiva (v.8.0.1) software and analyzed using FlowJo v.10.7.2 (Tree Star).

### Droplet-based single-cell sequencing

Sorted single cells were captured using Chromium controller (10x Genomics) according to the Chromium Single Cell 5’ Reagent Kit (CG000330 Rev C) protocols. Briefly, the cell suspension was loaded onto the controller to encapsulate single cells into droplets with barcoded gel beads using Gel Bead kit v2 (10X Genomics, 1000265) and Next GEM Chip K Single-Cell kit (10X Genomics, 1000286). Up to 20,000 cells were added to each channel with expected recovery of 8,000 cells. Captured cells were lysed and the released RNA was barcoded through reverse transcription. The 5ʹ gene expression (GEX) libraries, V(D)J libraries (10x Genomics, 1000190) and cell surface protein libraries (10X Genomics, 1000256) were prepared according to manufacturer’s protocols. Library quality was assessed using a 2200 TapeStation (Agilent). Libraries were sequenced on an Illumina NovaSeq6000 platform (150+150 bp paired read length).

### scRNA-seq processing and analysis

Libraries were demultiplexed using bcl-convert v3.9.3. Reads were mapped to human genome (GRCh38-2020-A) using 10x Genomics Cell Ranger v6.1.2 multi. Hashtag-based sample demultiplexing was done using hashedDrops function from DropletUtils R package v1.19.3. Theollowing cells were filtered out: no confident sample assignment or classified as doublet based on the hashtag read count, >1% of hemoglobin gene expression, outliers in the distribution of mitochondrial gene expression or the number of detected genes. Outliers were detected based on the median absolute deviation using isOutlier function from scuttle package v1.9.4. Immunoglobulin gene counts were summed in one feature prior to normalization, as well as ribosomal protein genes and HLA genes. Data normalization and variable gene selection was done using SCTransform from Seurat package v4.3.0. Principal component analysis, UMAP embedding, nearest neighbor graph construction and clustering were done using standard Seurat functions. Clusters of cells expressing non-B cell markers (*LYZ, CD14, CD68, GNLY, GZMA, CD3E, CD3G, CD4*) were removed and the data was re-normalized and re-clustered. Diffusion map embeddings were calculated using destiny R package v3.8.1. Single-cell pseudotime trajectory was calculated with slingshot R package v2.7.0 based on the clusters defined with Seurat. S-reactive cells were defined based on having read count above the threshold for either Spike or RBD antigen barcode. Thresholds were defined based on the optimal separation of positive and negative cell populations.

### Gene expression and signature enrichment analysis

Differentially expressed genes were detected using FindMarkers with Benjamini-Hochberg multiple testing correction. The genes were considered differentially expressed if p.adjusted <0.05 and average fold change >1.3. Ingenuity Pathway Analysis (IPA) was performed using avg_logFC values of all differentially expressed genes. BCR signaling pathway expression was quantified using AUCell v1.21.1.

### BCR repertoire analysis

Full-length V(D)J contigs were assembled with Cell Ranger and aligned to the IMGT reference using IgBLAST v1.20.0 and all downstream analyses were done with R. Contigs were filtered using the following criteria: cells passed the transcriptome quality control, V(D)J information is available for one heavy and one light chain, classified as full-length, productive and high-confidence by Cell Ranger. Clones were defined as groups of cells sharing at least one V and J genes in the top-3 alignment hits and having CDR3 nucleotide sequence similarity >80%. The similarity threshold was defined based on the distribution of nearest-neighbor similarity. After defining clones, all cells in a clone were assigned the V and J genes used by the majority of cells in a clone. To reconstruct phylogenetic trees, merged heavy and light chain V gene sequences were aligned using ClustalW method implemented in msa R package v1.31.7. Phylogenetic trees were reconstructed by RAxML v8 using germline sequence as an outgroup for each clone. Trees were visualized using ggtree v3.7.2. Low- and high-SHM cells were separated based on bimodal distribution of SHM counts^32^. SHM count threshold was set at a saddle point separating the first density peak from the rest of the distribution individually for each time point (Extended Data Fig. 2a).

### Recombinant monoclonal antibody production

Selected pairs of Ig heavy and light chain gene sequences were synthesized by Twist Bioscience and cloned into IgG1 and Igλ or Igκ expression vectors. Plasmids encoding paired Ig heavy and light chains were co-transfected into human embryonic kidney (HEK) 293F cells (Thermo Fisher Scientific) for recombinant mAb production. HEK293F cells were cultured in FreeStyle 293-F medium (Gibco) according to the manufacturer’s protocol.

### Enzyme-linked immunosorbent assay (ELISA)

ELISAs were performed as previously described^33^. In brief, 384-well high-binding polystyrene plates (Corning) were coated overnight at 4°C with Spike (4μg/ml), HCoV-HKU1 (4μg/ml), HCoV-OC43 (4μg/ml; Sino Biological), HCoV-229E (4μg/ml; Sino Biological), HCoV-NL63 (4μg/ml; Sino Biological), or RBD (4μg/ml) in Phosphate-buffered Saline (PBS). Plates were washed 3 times with washing buffer (1× PBS with 0.05% Tween-20 (Sigma-Aldrich)). ELISA plates were blocked with 4% bovine serum albumin (BSA) in PBS (serum ELISA), or 2% BSA in PBS (antigen ELISAs) for 1 hour at room temperature. Immediately after blocking, serially diluted serum samples at an initial dilution of 1:200 in 1% BSA with PBS, or mAbs at a starting concentration of 10 μg/ml (antigen ELISA), were loaded on the plate and incubated for 2 hours at room temperature. Plates were washed 3 times with washing buffer and then incubated with anti-human IgG-, IgA-, or IgM-horseradish peroxidase at 1:1000 (Jackson ImmunoResearch) in the corresponding blocking buffer (1× PBS with 0.05% Tween-20 and 0.01 M EDTA). Plates were developed by addition of the 1-Step ABTS substrate (Roche). mAbs S309^34^ and mGO53^35^ were used as positive and negative controls, respectively. Area under the curve (AUC) values were calculated using GraphPad Prism v9 (GraphPad) or R v4.3.1.

### Surface plasmon resonance (SPR)

SPR measurements were performed using a Biacore T200 (GE Health-care) instrument docked with Series S Sensor Chip CM5 (GE Healthcare), as previously described^36^. Briefly, anti-human Igκ and Igλ antibodies were immobilized on the chip using an amine coupling-based human antibody Fab capture kit. Hepes (10 mM) with 150 mM NaCl at pH 7.4 was used as a running buffer. Equal concentrations of the sample antibody and isotype control (mGO53)^35^ were captured in the sample and reference flow cells, respectively. Running buffer was injected at a rate of 10 μl/min for 20 min to stabilize the flow cells. RBD at 0.02, 0.08, 0.31, 1.25 and 5 μM in running buffer was injected at a rate of 30 μl/min. The flow cells were regenerated with 3 M MgCl2. Steady-state dissociation constants were calculated using BIACORE T200 software V2.0.

### Statistical analysis

Statistical analyses were performed using R v4.3.1 or Prism v9 (GraphPad) using tests described in the figure legends. All experiments were performed at least in duplicate.

## Data availability

The raw and processed sequence data have been deposited in GEO under the accession number GSE244297.

## Code availability

The code for transcriptome and BCR analysis is available on Github (https://github.com/obrzts/SARS-CoV-2_vac_scRNAseq).

## Author contributions

H.W. conceived the study. Z.L., A.O., R.M. and H.W. designed the experiments. K.B. collected blood samples. J.M. and R.M. isolated PBMCs and serum. M.v.S. and E.S. provided Spike and RBD proteins. Z.L. and F.S. sorted single cells and prepared sequencing libraries. A.O. performed computational analyses. Z.L. expressed monoclonal antibodies. Z.L., O.E.O. and J.M. performed serum and monoclonal antibody assays. Z.L., A.O. and H.W. interpreted the data. Z.L., A.O. and H.W. wrote the paper. H.W. acquired funding and supervised the study. All authors read and approved the final manuscript.

## Acknowledgments

The authors thank Yasmin Bergmann, Christine Niesik, Dorien Foster for technical assistance; Alessandro Greco and Thomas Höfer for advice on transcriptome analysis; the EMBL Protein Expression and Purification Core Facility, the DKFZ NGS Core Facility, DKFZ Single-Cell Open Lab (scOpenLab) and the DKFZ/EMBL/Heidelberg University Chemical Biology Core Facility, especially Peter Sehr, for technical assistance and services; Jan-Philipp Mallm for advice on single-cell RNA-seq; Julia Ludwig for helpful discussions. This work was supported by the Helmholtz Association’s Initiative and Networking Fund project “Virological and immunological determinants of COVID-19 pathogenesis—lessons to get prepared for future pandemics (KA1-Co-02_CoViPa).” A.O. was supported by the DKFZ International PhD Program.

