## Extended Data Figures 1-4 for "Memory B cells dominate the early antibody-secreting cell response to SARS-CoV-2 mRNA vaccination in naïve individuals independently of their antibody affinity"

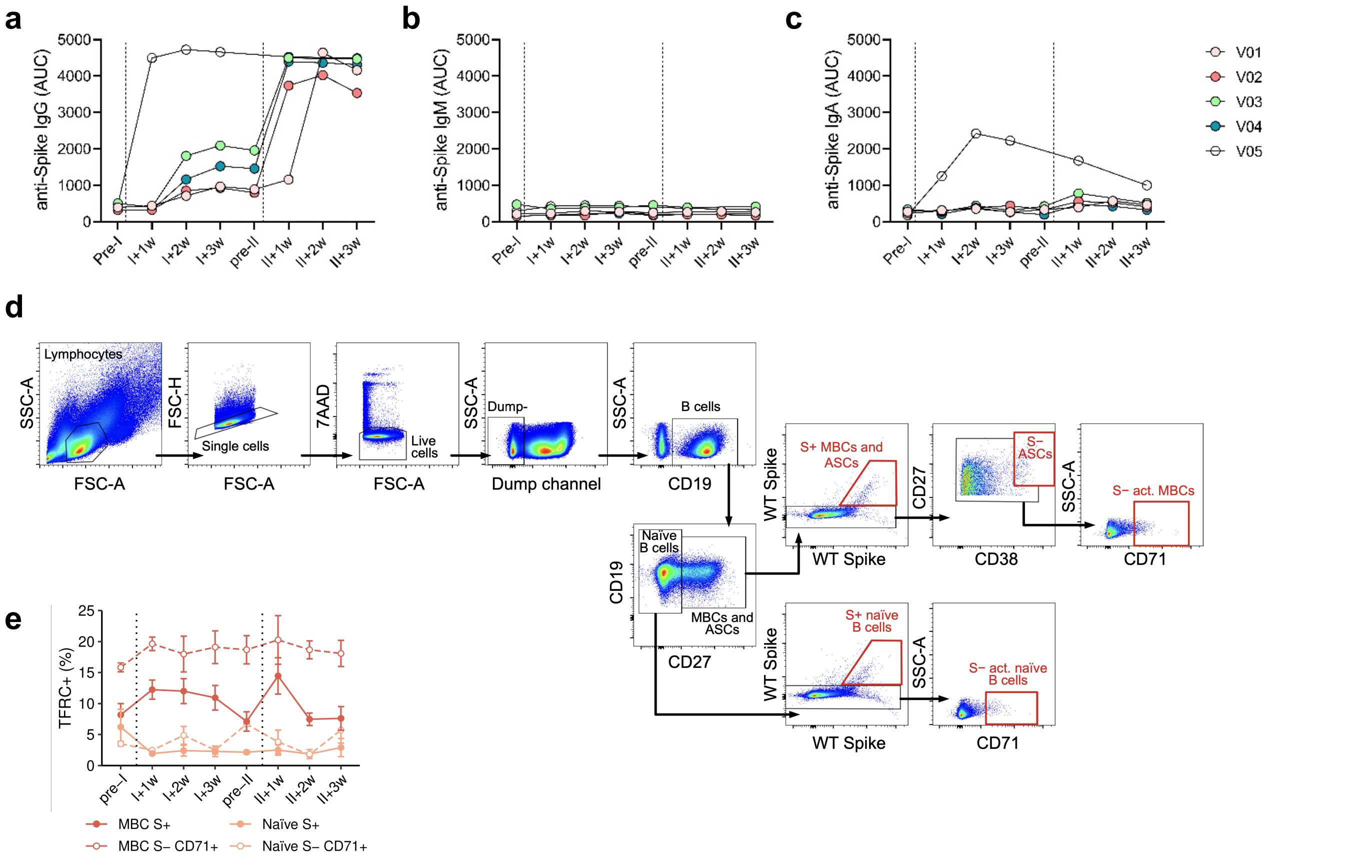


**Extended Data Fig. 1: Humoral and B cell anti-S response. a-c,** IgG, IgM and IgA anti-S serum response. Volunteer V5 rapidly developed anti-S IgG and IgA response indicating previous infection and shown as a positive control. **d,** Representative flow cytometric gating strategy for sorting 7AAD-CD3-CD8-CD14-CD16-CD19+ cells with S+CD27+ (S+ MBCs and ASCs), S+CD27- (S+ naïve B cells), S-CD27+CD38+ (S- ASCs) or S-CD71+ (S- activated naïve or MBCs). Red color marks the gates selected for sorting. **e,** Frequency of *TFRC*+ among MBCs and naïve B cells. *TFRC*+ cell were defined by presence of at least one *TFRC* transcript. Dotted vertical lines indicate prime and boost.

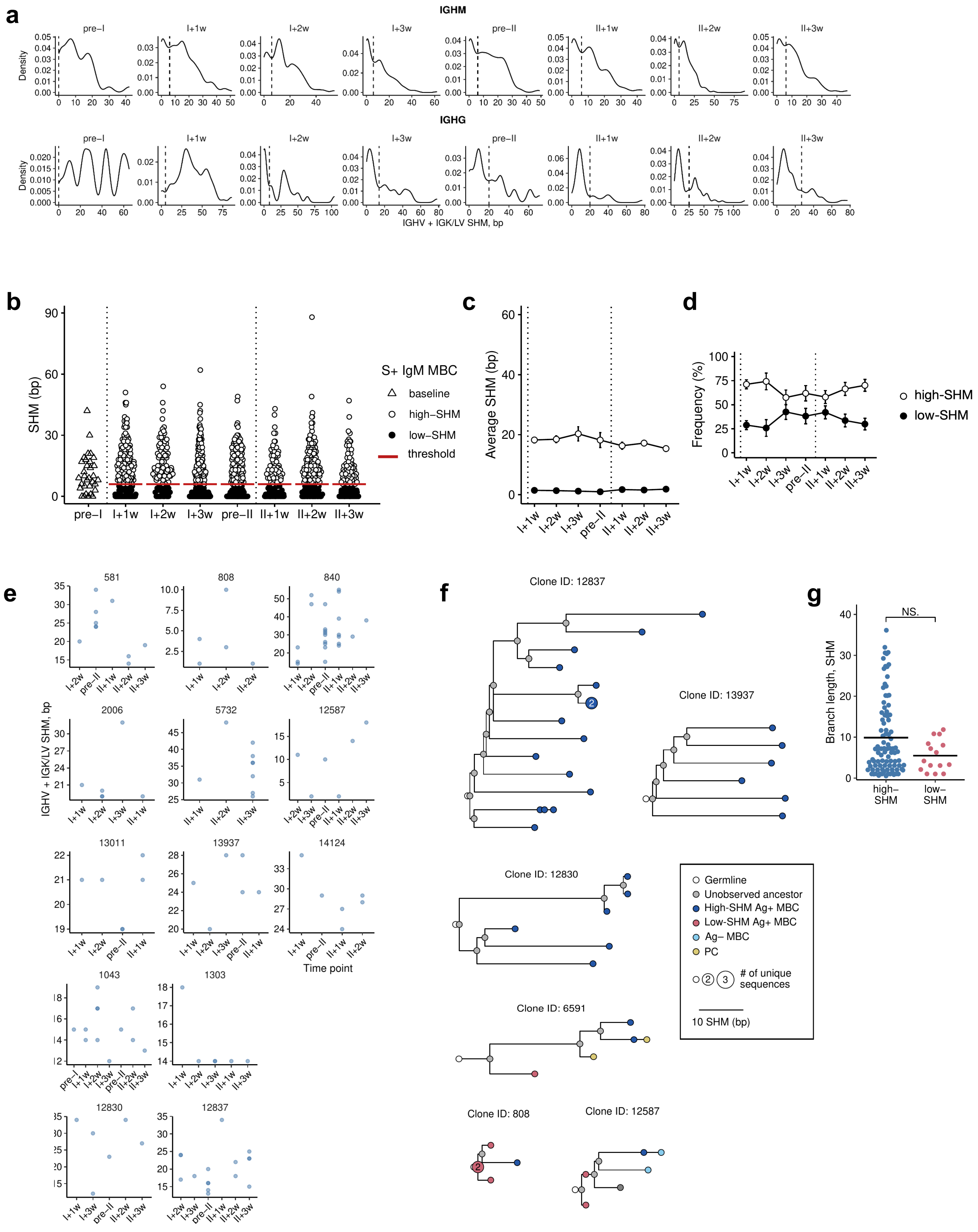


**Extended Data Fig. 2: BCR repertoire analysis of high- and low-SHM S+ MBCs. a,** SHM count distribution in IGHM and IGHG expressing S+ MBCs at the indicated time points. Dashed lines show thresholds separating the first density peak from the rest of the distribution. **b-d,** S+ IgM MBC response. *IGHV + IGK/LV* SHM counts (**b**), average SHM counts over time (**c**) and frequency of low- and high-SHM cells over time (**d**). Dotted vertical lines indicate prime and boost. **e,** SHM counts in the high-SHM MBC clones with at least 5 members and detected at at least 3 time points. **f,** Phylogenetic trees of three largest MBC clones in high-SHM (top) and low-SHM (bottom) S+ MBCs. Branch length represents number of somatic mutations in V genes of heavy and light chain, circle (node) size corresponds to the number of sequences also indicated as a number in circle. Germline sequence is shown in white, unobserved ancestor sequences in grey, observed sequences are colored. **g,** Branch length distribution in high- and low-SHM lineage trees. NS. - not significant, two-tailed Mann-Whitney test.

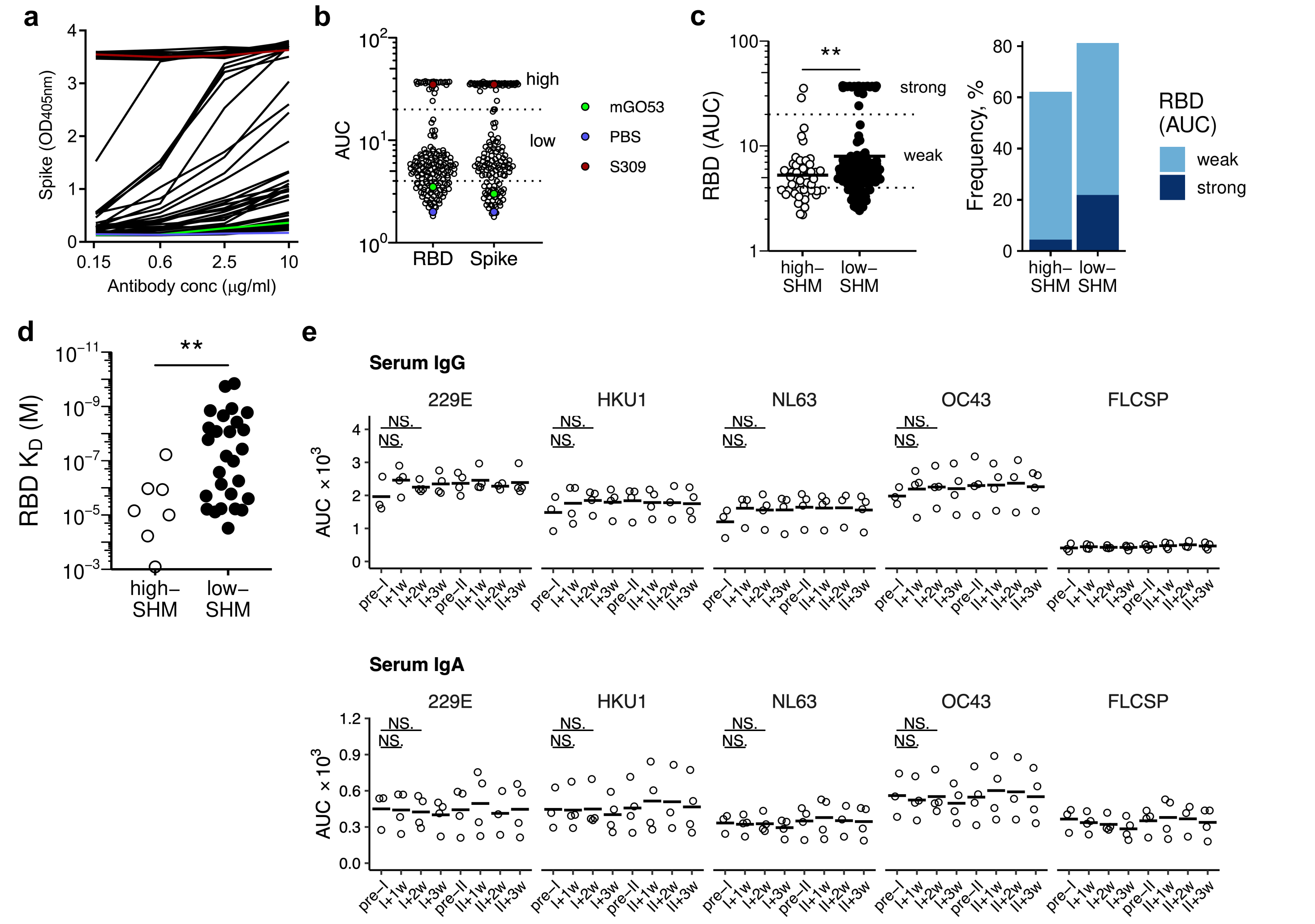


**Extended Data Fig. 3: Antigen binding of monoclonal and serum antibodies**. **a-b,** ELISA-reactivity to Spike and RBD. Representative optical density (OD; n=64) (**a**) and area under curve (AUC) distribution and thresholds for non-, weak- and strong-binding mAbs (**b**; n=141). mAb mGO53 and PBS were used as negative controls and mAb S309 as a positive control. **c,** AUC values from RBD ELISA (left) and frequency of RBD binders (right) in high- and low-SHM S+ MBCs. **d,** RBD binding affinity of mAbs from high- and low-SHM S+ IgG MBCs measured by SPR (non-binders excluded). **e,** Serum antibody response against HCoV. FLCSP (full-length circumsporozoite protein from *Plasmodium falciparum*) was used as a negative control. Open circle dots represent individual donors. Data are representative of at least two independent experiments. All antibodies were cloned and expressed as IgG1. NS. – not significant, *P < 0.05, **P < 0.01, two tailed Mann-Whitney test.

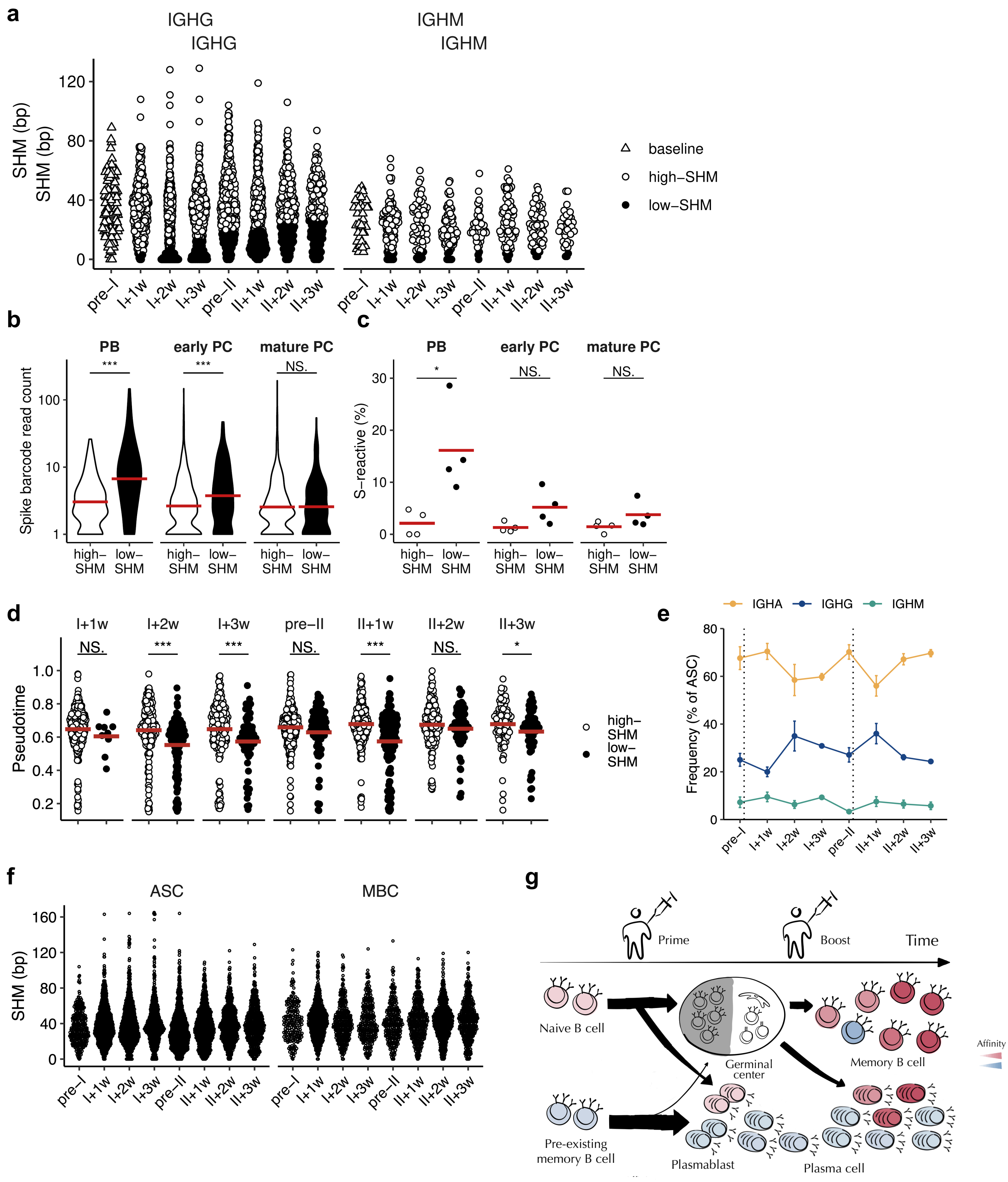


**Extended Data Fig. 4: Ig gene repertoire, antigen binding and transcriptome analysis of ASCs. a,** *IGHV + IGK/LV* SHM counts. **b,** Spike barcode read count distribution in high- and low-SHM ASCs. PB, early PC and mature PC are ASCs. IgM and IgG ASCs are pooled. **c,** Frequency of S+ cells in subpopulations of ASCs. PB, early PC and mature PC are ASCs. **d,** Pseudotime values of low- and high-SHM ASCs over time. **e,** Antibody isotype frequencies of ASCs over time. **f,** *IGHV + IGK/LV* SHM counts in IgA MBCs and ASCs. **g,** Proposed biological model. *P < 0.05, **P < 0.01, ***P < 0.001, ****P < 0.0001, NS, not significant; two tailed Mann-Whitney test.
